# Suppress me if you can: neurofeedback of the readiness potential

**DOI:** 10.1101/2020.08.07.241307

**Authors:** Matthias Schultze-Kraft, Vincent Jonany, Thomas Samuel Binns, Joram Soch, Benjamin Blankertz, John-Dylan Haynes

## Abstract

Voluntary movements are usually preceded by a slow, negative-going brain signal over motor areas, the so-called readiness potential (RP). To date, the exact nature and causal role of the RP in movement preparation have remained heavily debated. One important open question is whether people can exert conscious control over their RP, for example by learning to suppress it. If people were able to initiate spontaneous movements without eliciting an RP, this would challenge the idea that the RP is a necessary stage of the causal chain leading up to a voluntary movement. We tested the ability of participants to control the magnitude of their RP in a neurofeedback experiment. Participants performed self-initiated movements and after every movement they were provided with immediate feedback about the magnitude of their RP. They were asked to find a mental strategy to perform voluntary movements such that the RPs were as small as possible. We found no evidence that participants were able to to willfully modulate or suppress their RPs while still eliciting voluntary movements. This suggests that the RP might be an involuntary component of voluntary action over which people cannot exert conscious control.

## 1. Introduction

The readiness potential (RP) is a slow scalp negativity observed over motor areas in the electroencephalogram (EEG). It can start more than one second before spontaneous, voluntary movements (Kornhuber & Deecke, 1965; Shibasaki & Hallett, 2006). One traditional account of the RP is that it is a causal precursor to voluntary action and that it reflects an unconscious decision to act (Libet et al., 1983; Libet 1985). While recent studies indeed suggest that the RP is involved in the formation of conscious intention (Pares-Pujolras et al., 2019; Schultze-Kraft et al., 2020) and that it is a signal specific to voluntary action (Travers et al., 2020), other studies have raised questions about its role in movement preparation (Schurger et al., 2012; Schmidt et al., 2016; Schurger, 2018) and its role in human volition has remained unclear (Frith & Haggard, 2018).

The precise causal role of the RP in movement preparation notwithstanding, it is frequently assumed that it is a necessary part of the causal chain that allows for voluntary action (although this is debated, see e.g. Radder & Meynen 2012). A related and more specific possibility could be that the RP is an “*involuntary* component of *voluntary* action”. That is, that the RP occurs automatically and irreversibly (i.e. involuntarily) once a person has voluntarily decided to move. In contrast, an alternative possibility is that people can exert conscious control over their RP, for example by learning to suppress or abolish it completely, while still being able to elicit spontaneous movements. If this were the case, it is conceivable for people to be able to execute voluntary movements that are not preceded by RPs. This possibility has not yet been tested directly.

In a recent study, participants executed self-paced movements and were occasionally interrupted by stop signals that were triggered by the detection of emerging RPs in the ongoing EEG (Schultze-Kraft et al., 2016). In one case they were instructed to try and “move unpredictably” so as to not cause an RP and thus in turn avoid causing stop signals. We found no evidence that participants attenuated their RP, which was the same across all conditions studied, despite the fact that people could have in theory increased their performance by suppressing their RP. However, it still remains unclear whether participants can decrease their RP when explicitly requested to do so. One way would be to provide people with immediate and graded neural feedback about the size of the RP they just produced and ask them to reduce it. This would be akin to an ‘autocerebroscope’ (Feigl, 1957, p. 456), a device that allows one to observe one’s own brain signals as one behaves. Such an approach could potentially enable people in a trial-and-error fashion to learn how to modulate and suppress their RPs, as with examples of neurofeedback for other cognitive processes (Papo, 2019).

We distinguish two principles which could enable people to achieve control over their RP. First, the readiness potential has been shown to be modulated by various attributes of voluntary movement, such as its inertial load and force deployment (Becker & Kristeva 1980; Kristeva et al., 1990; Slobounov et al., 2004), its complexity (Benecke et al., 1985; Simonetta et al., 1991; Kitamura et al., 1993), its purposiveness and selection mode (Masaki et al., 1998; Praamstra et al., 1995), and by explicit demands on timing (Bortoletto & Cunnington, 2010; Baker et al., 2012; Verleger et al., 2016). Further, compared to RPs observed in classical Libet-style studies (Libet et al., 1983), RPs are considerably smaller when spontaneous movements are executed unconsciously (Keller & Heckhausen, 1990), and almost absent when movements are initiated by deliberate, value-based decisions (Maoz et al., 2019). In all these studies, the modulation of the RP resulted from an experimental manipulation, that is by instructing participants to change specific characteristics of voluntary movements. Yet, it seems plausible that, when provided with trial-by-trial feedback of their RP, people would be able to identify how changing specific movement features allows them to modulate their RPs.

Second, studies have investigated the self-regulation of slow cortical potentials (SCP), which are polarizations of EEG that can last up to several seconds (Birbaumer 1999), and of which RPs are considered a specific type. Using a training based on visual feedback of SCP shifts and operant learning principles (Elbert et al., 1980; Rockstroh et al., 1984), people can learn to self-regulate their SCPs, which has been used in communication systems for paralyzed patients (Kübler et al., 1999; Kübler et al., 2001; Neumann et al., 2004). The mechanisms that allow such self-regulation are not well understood but are assumed to be based on a redistribution of attentional resources (Birbaumer 1999). This learning of self-regulation could in principle be employed by participants when provided with a trial-by-trial feedback of RP magnitude.

Here, we tested the possibility of a voluntary suppression of readiness potentials in a neurofeedback experiment. Our core research question was to test whether people could suppress RPs by purely mental efforts, and not by changing physical movement characteristics that are known to modulate RPs. Participants performed self-paced pedal presses in single trials. After each pedal press we used a machine learning approach to derive a score that reflected the size of the RP that had just been produced and that was shown to participants as feedback. Participants were challenged to find a mental strategy to perform movements such that the scores (and therefore their RPs) were as small as possible.

## 2. Methods

### 2.1. Participants

Based on the average sample size of previous studies (Schultze-Kraft et al., 2016; Schultze-Kraft et al., 2020; Pares-Pujolras et al., 2019; Schurger et al., 2012), we aimed for a minimum sample size of 15 participants. Considering that some would have to be excluded, we tested a total of 22 participants. Following our exclusion criteria (see below), 19 participants were included in the final sample. (11 female, mean age 26.9, SD 5.7 years). The experiment was approved by the local ethics board and was conducted in accordance with the Declaration of Helsinki. All participants gave their informed oral and written consent, and were paid €10 per hour.

### 2.2. Experimental setup

Participants were seated in a chair facing a computer screen at a distance of approximately 1 m. They were asked to place their hands in their lap and to position their right foot to the right of a 10 cm x 20 cm floor-mounted switch pedal (Marquardt Mechatronik GmbH, Rietheim-Weilheim, Germany). Throughout the experiment, EEG was recorded at 1 kHz with a 64-electrode Ag/AgCl cap (EasyCap, Brain Products GmbH, Gilching, Germany) mounted according to the 10-20 system and referenced to FCz and re-referenced offline to a common reference. EEG was recorded from the following 51 electrodes: AF7, AF3, Fpz, AF4, AF8, FT7, F5, F3, F1, Fz, F2, F4, F6, FT8, FC5, FC3, FC1, FC2, FC4, FC6, C5, C3, C1, Cz, C2, C4, C6, CP5, CP3, CP1, CPz, CP2, CP4, CP6, TP7, P5, P3, P1, Pz, P2, P4, P6, TP8, PO3, PO1, POz, PO2, PO2, O1, Oz, O2. In order to obtain the earliest measure of movement onset, 3D acceleration of the right leg was recorded with an accelerometer (Brain Products GmbH, Gilching, Germany) that was attached with an elastic band to the right calf. The amplified signal (analog filters: 0.1, 250 Hz) was converted to digital (BrainAmp MR Plus and BrainAmp ExG, Brain Products GmbH, Gilching, Germany), saved for offline analysis, and simultaneously processed online by the Berlin Brain-Computer Interface toolbox (BBCI, github.com/bbci/bbci_public). The Pythonic Feedback Framework (Venthur et al., 2010) was used to generate visual feedback.

### 2.3. Experimental design

The experiment consisted of two stages (Fig. 1), a *preparatory* stage, and a *feedback* stage. The preparatory stage was performed to obtain data for training a classifier in preparation for the subsequent feedback stage. During the preparatory stage participants performed a simple self-paced movement task. The start of a trial was signaled by a white circle appearing on the screen. Participants were instructed to wait for roughly 2 seconds, after which they could press the pedal at any time. In accordance with standard definitions of the readiness potential they were asked to avoid pre-planning the movement, avoid any obvious rhythm, and to press when they felt the spontaneous urge to move (Kornhuber & Deecke, 1964; Libet et al., 1983). When the pedal was pressed the white circle turned red for 1 second, after which it disappeared and was replaced by a fixation cross. This constituted the end of a trial. The fixation cross remained onscreen for a 3 s intertrial period. Each participant performed a total of 100 trials in the preparatory stage, with the possibility of taking a break after each 25 trials.

**Fig. 1.**
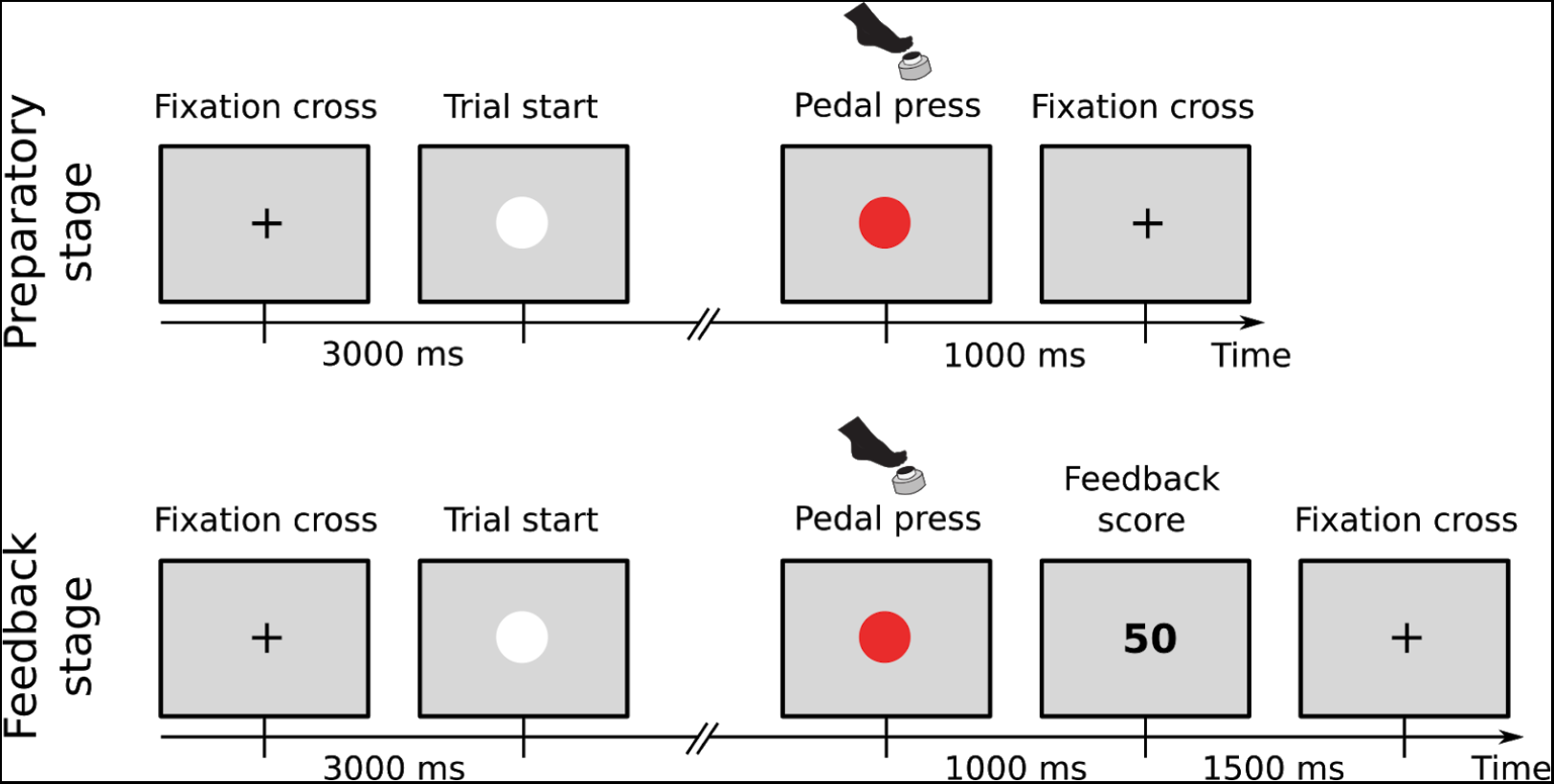
Experiment paradigm. In both the preparatory and the feedback stage, trial start was signaled by a white circle appearing on the screen. When a pedal press was executed, the circle turned red for 1 s. In the preparatory stage, the trial ended and a fixation cross was shown for an intertrial period of 3 s. In the feedback stage, before the trial ended a number was shown on the screen for 1.5 s, after which the fixation cross was shown.

During the second part of the experiment, the feedback stage, participants again performed self-paced pedal presses in single trials, as during the preparatory stage. However, after the participants had moved an integer number was displayed on the screen for 1.5 s. Participants were informed that “this number reflects a brain signal recorded when you decided to press the pedal. Larger numbers mean large signals, small numbers mean small signals”. They were given the additional task to develop strategies to achieve preferably low numbers. Participants were (i) instructed to move spontaneously and to not execute abnormal (e.g. very slow, interrupted) movements, but were otherwise free to find a strategy to achieve the goal, (ii) informed that, based on noisy measurements, scores might greatly vary from trial to trial and that they might thus need many trials to realize if a strategy works or not, and (iii) instructed to keep using a strategy to further lower the scores if they happen to find one that works. Participants performed 300 trials during the feedback stage, with the possibility of taking a break after each 25 trials.

### 2.4. Training of classifiers from preparatory stage data

Before the feedback stage, we performed three consecutive analyses on the data recorded during the preparatory stage: (1) We trained an *accelerometer classifier* that detected physical movement onset times in realtime from accelerometer data, (2) we selected the most informative EEG channels from the independent preparatory data, and (3) we trained a real-time *EEG classifier*. Both the real-time accelerometer and EEG classifiers were then used during the feedback stage to assess the movement and the RP produced in each trial and to derive a score in real-time that was shown to participants as feedback at the end of the trial.

#### 2.4.1. Detection of movement onsets from accelerometer

The accelerometer device attached to the right calf recorded acceleration in the direction of three orthogonal space axes. We determined the time of movement onset in each trial with a variance-based approach. We trained a linear classifier on log-variance features extracted from two time windows: (1) a time window from −200 to 0 ms, time-locked to pedal press (“movement” class), and (2) a time window from 300 to 500 ms, time-locked to trial start (“idle” class). The former time window was expected to contain the acceleration of the foot during the movement, and thus have a large variance, while in the latter the acceleration was expected to be at baseline during the instructed self-paced waiting time of 2 s. In order to determine the movement onsets of each trial, a classifier was trained on the “movement” and “idle” time windows of 99 trials, and then applied with a sliding window on the remaining trial. The analysis worked backward from the physical completion of the pedal press, looking for the last time window preceding the pedal press where there was no evidence for movement. For this, a first window was time-locked to the pedal press and then it was sample-wise shifted back in time until the classifier output indicated being in the “idle” class. The time of this last idle window before movement was registered as the time of movement onset. This procedure was applied to each of the 100 trials per participant in a leave-one-out scheme. Trials with movement onsets times 3 standard deviations below or above the individual mean were excluded from further analysis. Finally, a classifier (hereafter referred to as “accelerometer classifier”) was trained on the accelerometer data from all remaining trials and subsequently used during the feedback stage for real-time detection of movement onset (see below).

#### 2.4.2. EEG channel selection

We preselected a subset of channels that would be used for the assessment of RP magnitude during the feedback stage. This selection was done using the independent data recorded during the preparatory stage. By selecting channels near the vertex, we focused on channels where the RP is assumed to predominate, and further aimed to minimize the impact of movement or eye artifacts that predominantly occur at peripheral electrodes. For each of the selected trials of the preparatory stage, we defined two EEG segments: (1) a 1000 ms long segment time-locked to and preceding movement onset (“movement” class), and (2) a 1000 ms long segment time-locked to and preceding trial start (“idle” class). The former were expected to contain an RP-typical negativation of EEG signals at certain channels, while the latter did not contain RPs. For each segment, we subtracted the average signal in the last 200 ms of the segment from the average signal in the first 200 ms of the segment. For each segment, this value thus represented how much the signal had changed in the 1000 ms preceding either movement onset, or trial start, respectively. For each EEG-channel individually, we then performed two one-sided t-tests in order to test (1) if the signal changes throughout the segment in the “movement” class were smaller than zero (to reflect the negative-going RP), and (2) if the signal in the “movement” class was smaller than that in the “idle” class (to account for potential negative-going signal drifts in the idle condition). The criterion for selecting a channel was then that the null hypothesis of both these tests on the preparatory data could be rejected at an alpha level of 0.05. The number of selected channels thus varied between participants. Fig. 2 shows a topographic heatmap of the frequency with which channels were selected across participants. Channel Cz was selected for all 22 participants, reflecting the fact that readiness potentials preceding foot movements are typically most distinct over that channel (Brunia et al., 1985; Schultze-Kraft et al., 2016). Channels further away from Cz were selected with less frequency. On average, 10 (SEM=1) channels were selected per participant.

**Fig. 2:**
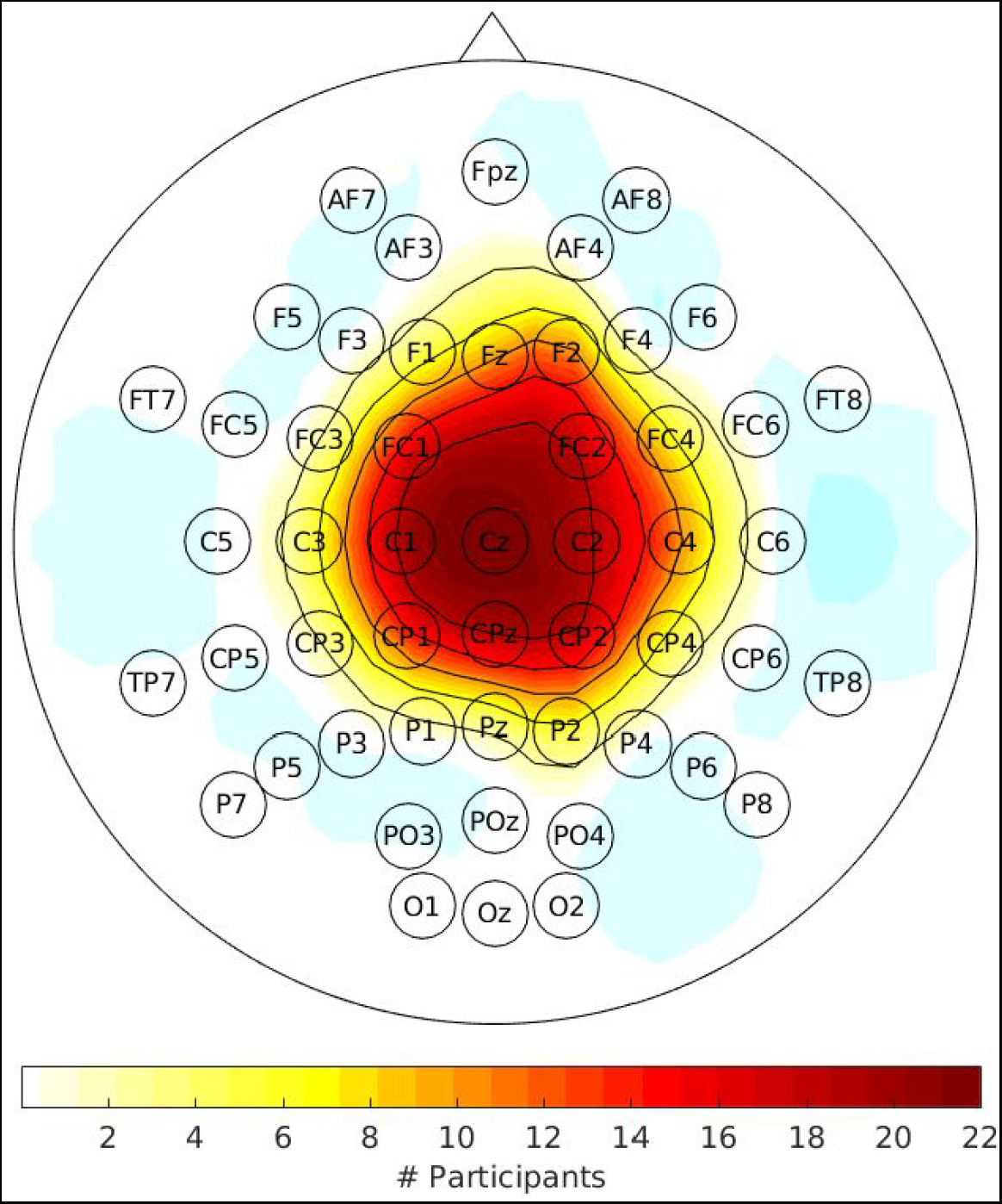
Topography of selected channels. The scalp topography shows a heat map of the number of times each channel was selected across participants.

#### 2.4.3. Training of EEG classifier

In order to extract RP-related spatio-temporal features from the EEG, we performed the following analysis, using data from the preparatory stage: For each trial and for each of the selected channels, we defined two EEG segments: (1) a 1000 ms long segment time-locked to and preceding movement onset (“movement” class), and (2) a 1000 ms long segment time-locked to and preceding trial start (“idle” class). These segments were first baseline corrected in the interval −1000 to −900 ms and then downsampled by averaging the data in consecutive 100 ms intervals, thus obtaining 10 temporal features per segment and channel. Finally, these features were concatenated across all selected channels to obtain a spatio-temporal feature vector per segment. In order to derive an estimate of the distribution of classifier outputs for EEG segments containing RPs, we performed the following analysis: A regularized Linear Discriminant Analysis (LDA) classifier with automatic shrinkage (Blankertz et al., 2011) was trained on the “movement” and “idle” segments of all but one trial in the preparatory data, and then applied to the “movement” segment of the left out trial. This procedure was applied to each trial in a leave-one-out scheme, resulting in one classifier output value per single-trial RP. The mean μ_0_ and standard deviation σ_0_ of the resulting distribution were calculated. These values were used during the feedback stage for transforming the EEG classifier outputs into a feedback score (see below). Finally, the same classifier (hereafter referred to as “EEG classifier”) was trained on all trials and subsequently used during the feedback stage (see below).

### 2.5. Real-time feedback

During the feedback stage, every 20 ms both previously trained classifiers were applied to the real-time data acquired at that moment. That is, the accelerometer data acquired in the last 200 ms was subjected to the accelerometer classifier, and the EEG data acquired in the last 1000 ms was subjected to the EEG classifier. This yielded one output value per classifier at each sample point. The logic was as follows. First, we wait until the button is pressed. Then we use the accelerometer classifier to look back in time from the button press and identify the time of movement onset, defined as the classifier switching from “idle” to “movement” class. Then we extract the EEG classifier output at that time. Subsequently, the EEG classifier output value at the time of this movement onset was identified. Finally, to be easily interpreted by the participants, this value *x* was transformed to a score as

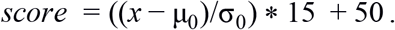

That is, after being normalized by the parameters obtained from the classifier outputs in the preparatory stage, the output was transformed such that an average value would result in a score of 50, and a value being one standard deviation above or below the mean would result in a score of 65 or 35, respectively. The resulting value was rounded to an integer and then showed to the participant as the feedback score after the pedal press.

### 2.6. Questionnaire

After finishing the feedback stage, participants were asked to fill out a questionnaire, which consisted of four questions: (1) “Overall, how much did you feel you could influence the scores shown on screen? (1 – not at all, 5 – a lot)”, (2) “How hard/easy was it to find a strategy that had an effect on the scores? (1 – very hard, 5 – very easy)”, (3) “Please use the table on the back of this sheet to write down your experience on the strategy/strategies that you used to achieve lower scores. On the left, please describe the strategy you used. On the right, please rate the success of the strategy and comment on anything that you find worth mentioning.”, and (4) “Did you have the feeling that one or more of the strategies work better over time, as if they were trainable? If so, which ones? (Please specify in the table)”.

### 2.7. Data selection

Before analysis, we performed a data selection approach based on two criteria.

#### 2.7.1. Accuracy of real-time movement onsets

We measured RPs in real-time by time-locking the EEG to the time of movement onset, not the time of the pedal press. As outlined above, our definition of movement onset uses the accelerometer classifier and looks backwards from the button press and identifies the latest time point before pedal press that is classified as idle (Fig. 3A). However, participants might not always perform smooth and continuous movements but instead perform multiphasic movements where they briefly pause or move slowly in between. In those cases, the accelerometer classifier at times failed to detect the true time of movement onset (Fig. 3B). Therefore, we had to ensure that movements onsets were not simply a later stage of a multiphasic movement with the participant having initiated the movement much earlier. Thus, we additionally required that there was no sign of movement in the phase before the detected movement onset. We excluded trials based on the following criterion: From trials in the preparatory stage, we defined the baseline variance of the accelerometer signals during rest. A feedback stage trial was then excluded if the accelerometer signal variance in the interval from −1000 to 0 ms before the real-time assigned movement onset was 3 SD above the baseline, thereby excluding on average 62 (SEM=15) trials per participant.

**Fig. 3:**
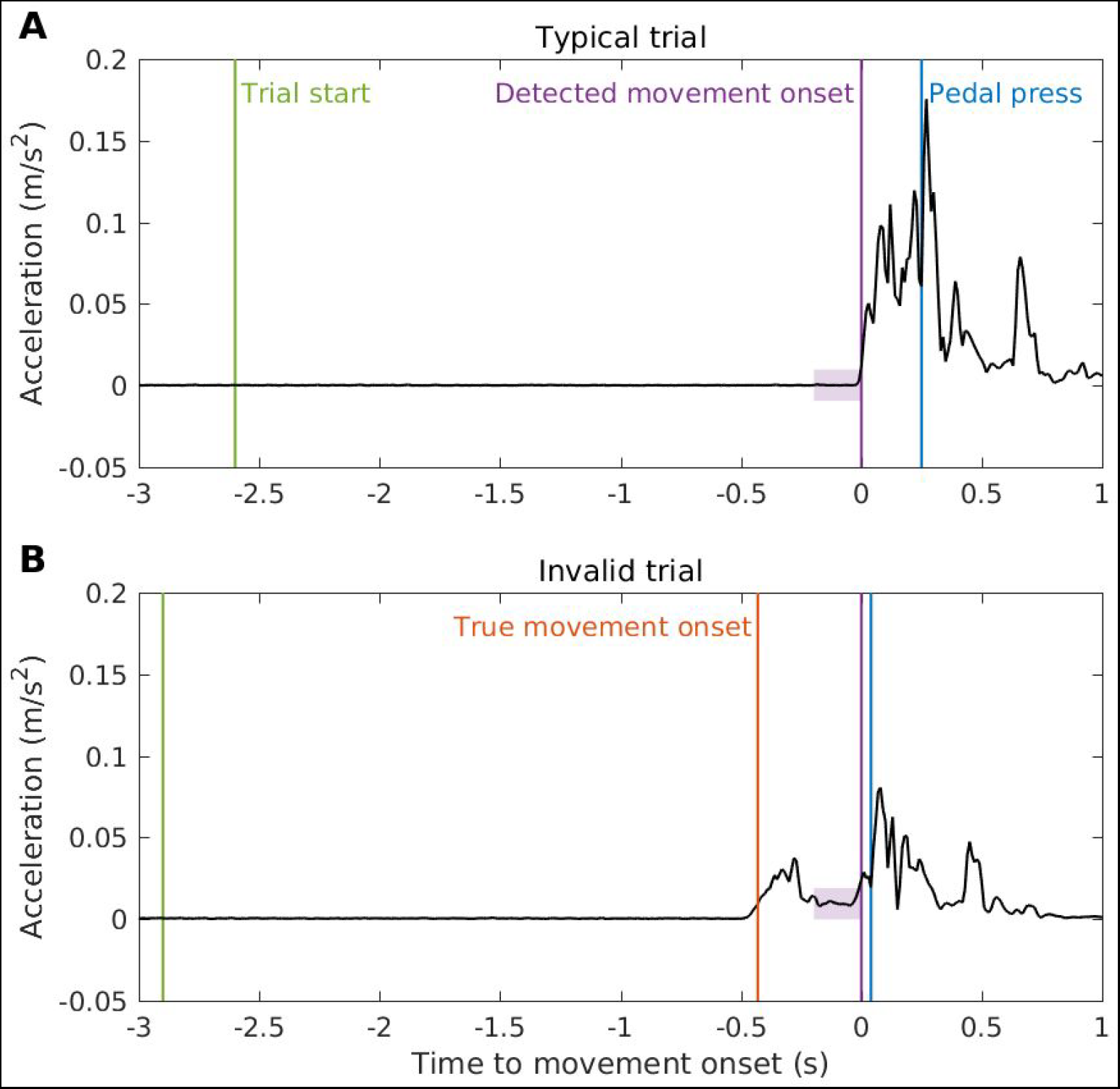
Exclusion of trials based on invalid movement onsets. Panels A and B show two exemplary single-trial traces of acceleration data from participant 20. Participants sometimes elicited interrupted, multiphasic movements (bottom) despite being instructed not to. So a selection procedure was designed to select trials with a single continuous movement by ruling out trials with a clear indication of movement *before* the movement onset detected in real-time. (**A**) Example of a valid movement trial. After trial start (green vertical line), the time of detected movement onset (purple vertical line, t=0 s) was defined as the last 200 ms window (shaded square) before pedal press (blue vertical line) in which the accelerometer classifier indicated the acceleration signal being in the idle class. (**B**) Example of an invalid movement trial. The backwards-looking algorithm detects an onset (defined here as time t=0 s) but the movement clearly started earlier. This is because after initiating the movement (red vertical line, t=-430 ms), the participant briefly slowed down the movement before pressing the pedal. The time point identified as movement onset by the accelerometer classifier was several 100 ms too late and at a physiologically impossible delay from the pedal press of 40 ms.

#### 2.7.2. Premature movement executions

We also focused on those trials where participants adhered to the instruction to wait for roughly 2 seconds after trial onset before deciding to press the pedal. This was to ensure that the time window used to extract RP features from the EEG (a 1000 ms window time-locked to and preceding movement onset) did not fall into the pre-trial start period. If participants did not follow this instruction, the extracted EEG features in that trial would be contaminated by the presentation of the trial start cue. Thus, we excluded trials where the delay between trial start and movement onset was less than 1000 ms, excluding on average 7 (SEM=5) trials per participant.

As Fig. 4 shows, the total number of trials excluded by these two criteria varied considerably across participants. Three participants with more than 50% excluded trials were excluded from all further analysis. The final sample thus included 19 participants, with an average of 255 (SEM=9) trials.

**Fig. 4:**
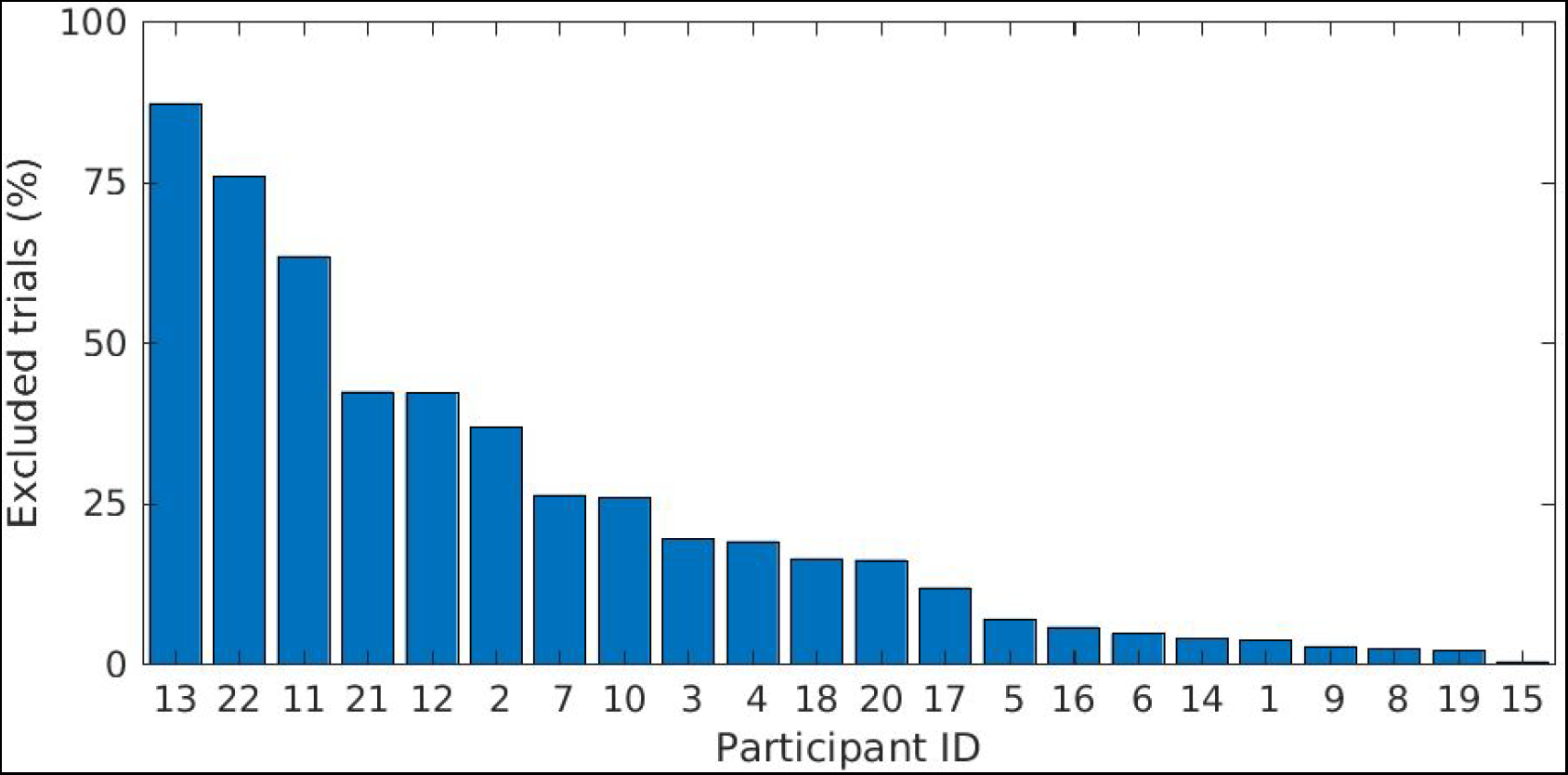
Percentage of excluded trials per participant. Participants are displayed in descending order according to the percent of trials excluded due to invalid online movement onsets or due to premature movement executions. Three participants (IDs 13, 22 and 11) with more than 50% excluded trials were excluded from further analysis.

### 2.8. Modelling feedback scores

Four explanatory variables were defined in order to examine the ability of participants to alter their RPs. One variable was trial number (*TN*), which was the key focus in this study: If participants were successful in gradually finding and training a strategy to lower their RP feedback scores during the *feedback* stage, this would be reflected in a decrease of RP feedback scores as a function of trial number. In addition, three additional measurements that characterize how participants generated the movement in each trial were defined as explanatory variables, despite not being the key focus here: Waiting time (*WT*, time from trial start to movement onset), movement duration (*MD*, time from movement onset to pedal press), and peak acceleration (*PA*, maximum acceleration measured between movement onset and pedal press). All four variables were z-transformed for each participant individually.

To test for an effect of the four variables on the RP, for each participant individually a linear regression was fitted on the trial-wise feedback scores (i.e. the linearly transformed EEG classifier outputs), using TN, WT, MD, PA and a constant regressor as predictors. This yielded one estimated regression coefficient for each participant and each variable, on which we then performed one-sample t-tests as a second-level analysis. Our main variable of interest was trial number: A gradual decrease of feedback scores in the course of the feedback stage would be reflected in a negative coefficient for the variable TN. Thus, a one-sided t-test was used to test whether the estimates were smaller than zero. For the movement characteristic variables WT, MD and PA, we had no specific assumption about the direction of the effect. Thus, for each of these variables, a two-sided t-test was performed. Finally, given the absence of an effect for all four variables (see Results), we validated the evidence for this absence using Bayesian t-tests, implemented in the open-source project JASP (Love et al., 2019). The prior used for the t-tests is described by a Cauchy distribution centred around zero and with a scale parameter of 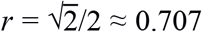, as suggested in Morey & Rouder (2011). Bayesian hypothesis testing aims to quantify the relative plausibility of the null and alternative hypotheses, and the Bayes Factor (BF) obtained by a Bayesian t-test is a continuous measure of evidence for either hypothesis (Keysers et al., 2020).

## 3. Results

### 3.1. Characteristics of movement execution

We first characterized the movements performed by participants during the feedback stage. As Fig. 5 shows, there was considerable between-subject variability in how movements were executed. Some participants initiated their movements on average after a couple of seconds after trial start, while others waited longer (mean waiting time 4119 ms, SEM 432 ms). Participants also required a different amount of time for the completion of the movement (mean movement duration 389 ms, SEM 38 ms), and with different peak acceleration (mean 7.79 cm/s^2^, SEM 0.71 cm/s^2^).

**Fig. 5:**
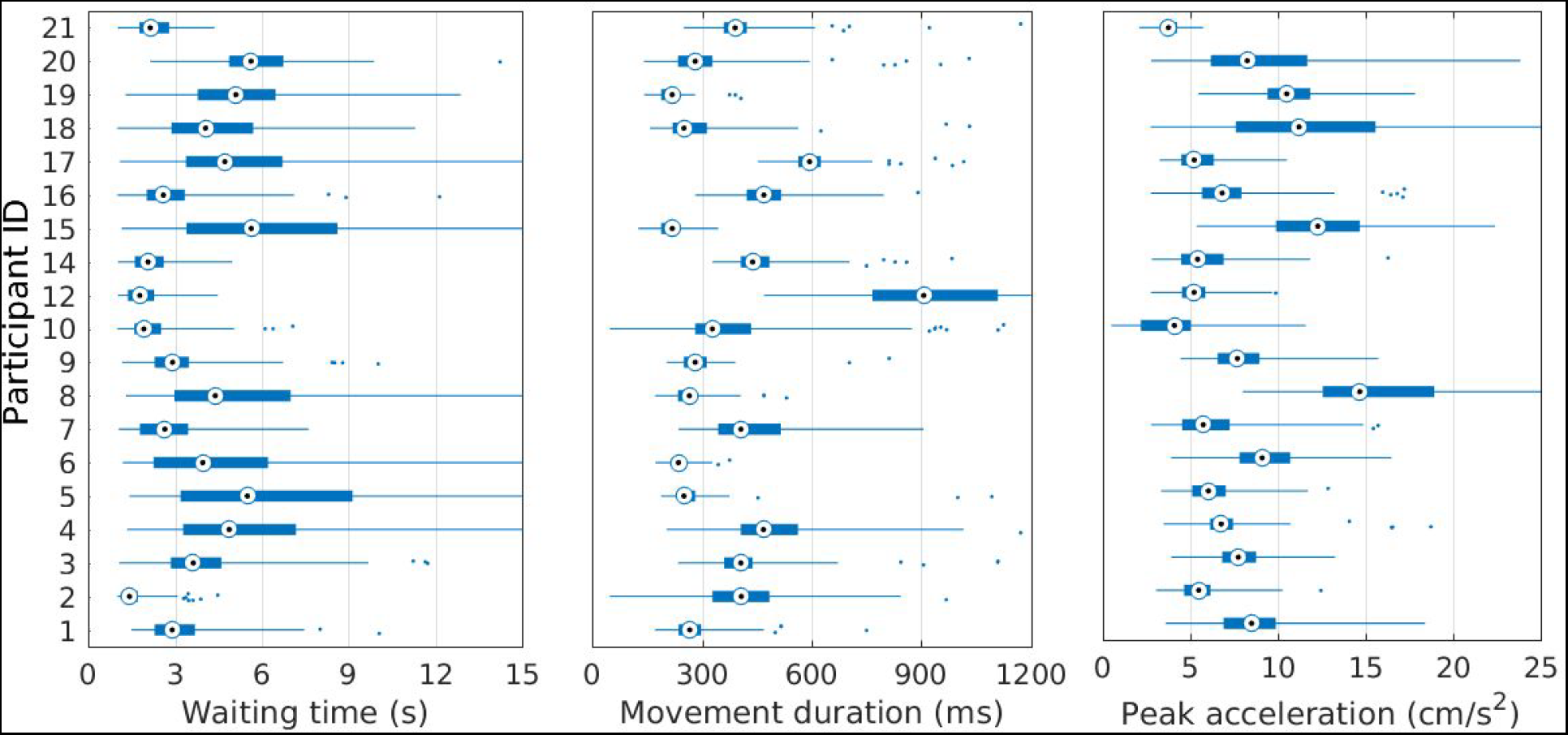
Movement characteristics during feedback stage. Boxplots show, for each participant individually, the distribution of waiting time, movement duration and peak acceleration of movements executed.

### 3.2. Validity of feedback scores

We checked that the feedback scores presented to participants indeed reflected the size of the RP (as would be expected by our method for defining feedback). Fig. 6A shows average RPs for the 5 different quintiles of feedback scores (low to high). RPs with high scores had early onsets and high amplitudes, whereas RPs with low scores had late onsets and small amplitudes. While RPs at all score levels had their largest amplitudes at the vertex (channel Cz), they were spatially less pronounced in more distant electrodes at lower scores. The correlation between feedback score and RP amplitude (Fig. 6B) was confirmed with a mixed-effects regression (*β*=−1.434, *p*<.001). Thus, on average, a decrease in 1.4 units of feedback score was equivalent to a 1 μV decrease in RP amplitude.

**Fig. 6.**
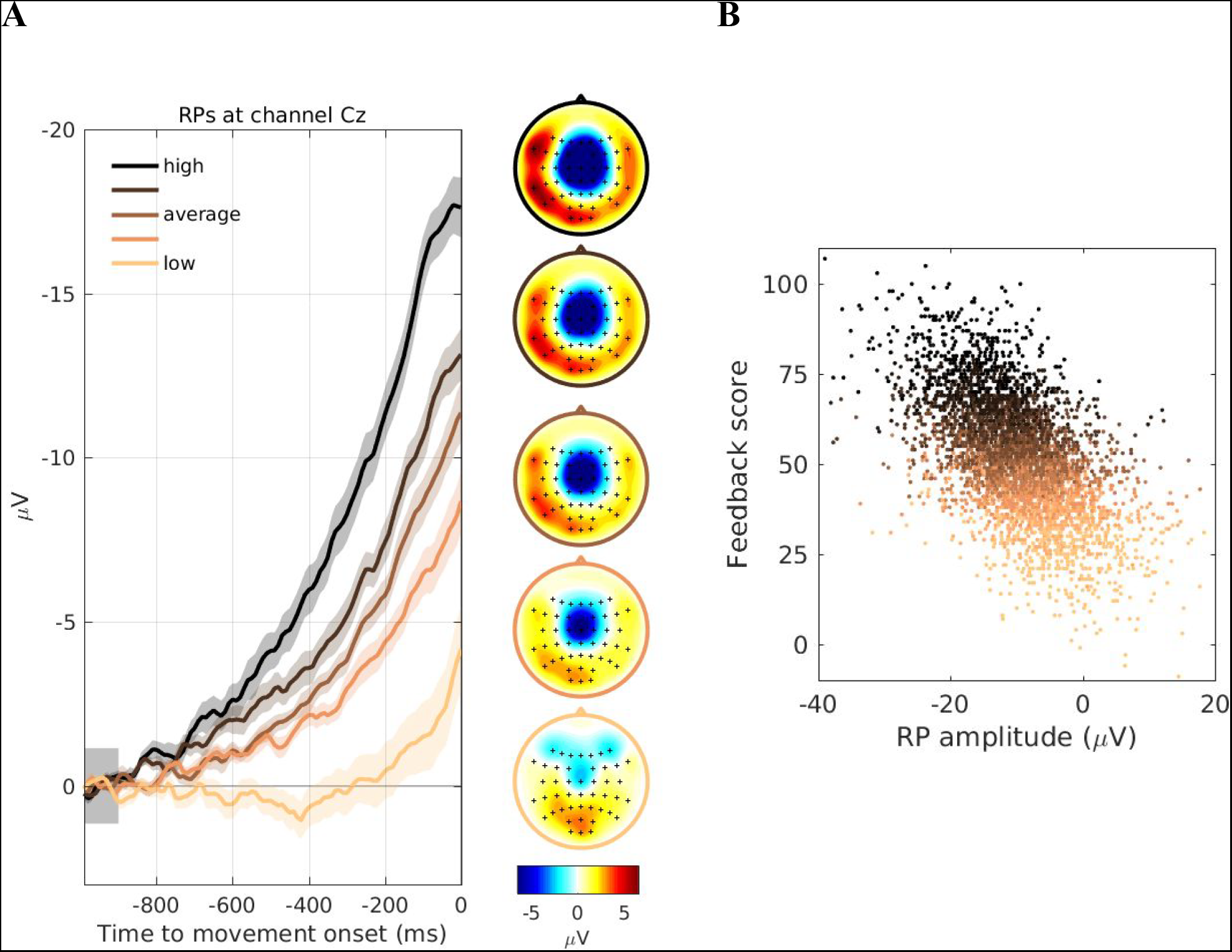
Waveforms, topographies and amplitudes of readiness potentials for different feedback levels. For each participant, trials were grouped into five quantiles depending on the feedback score that was calculated in real-time, color coded in all panels from low (light) to high (dark). (**A**) The left shows the grand average waveforms of the RPs at channel Cz for the five quintiles, baseline corrected in the interval [−1000 −900] ms. Standard error is shown as a shaded area. The right shows the corresponding scalp topographies of the average voltage in the interval [−100 0] ms for the five quintiles. (**B**) Amplitudes of single RPs, pooled across all participants and plotted against their corresponding real-time feedback score. Amplitudes were calculated as the average voltage at channel Cz in the time interval −100 to 0 ms. Please note that the RP is a negative brain signal so larger RPs correspond to “more left” values on the x-axis of the graph and are thus associated with higher feedback scores (hence the negative correlation). Taken together this sanity check confirms that the RP amplitudes assessed by the EEG classifier in realtime and fed back to the participants indeed reflected changes in RP waveform and amplitude.

### 3.3. Manipulation of scores by participants

We examined whether participants were successful in finding a strategy to execute movements with lower scores. If they were, this should be reflected in a gradual decrease of scores over the course of the 300 trials. A visual inspection of scores as a function of trial number showed no indication of such decrease (Fig. 7), and the shape RP waveform did not change over time (Fig. 8). A one-sided t-test on the regression coefficient estimates obtained for each participant (Fig. 9) showed that they were not smaller than zero (t_(18)_=0.103, p=0.541), and the Bayes Factor BF_0−_=4.539 indicates that the data are 4.5 times more likely under the null hypothesis which provides moderate evidence for absence (Jeffreys, 1961) of an effect of TN. The lack of a negative (linear) trend of feedback scores during the 300 trials of the feedback stage suggests that participants were not successful in finding a strategy to willfully reduce their RPs.

**Fig. 7.**
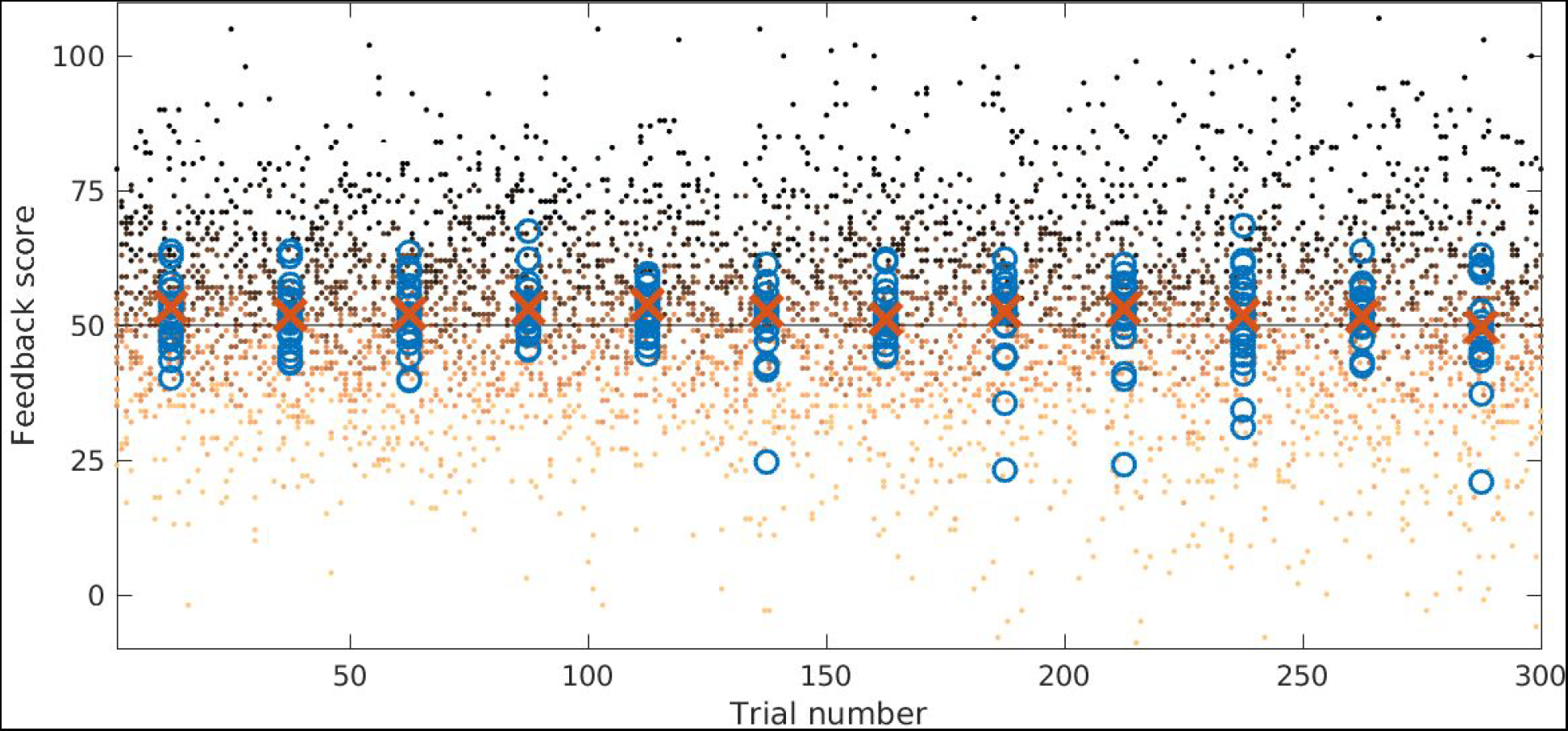
Change of feedback scores in the course of the feedback stage. Dots (color coded as in Fig. 6) show single-trial feedback scores, pooled across all participants, as a function of trial number. Blue circles show averages of individual participants, red crosses show population medians, both calculated over consecutive, non-overlapping bins of 25 trials.

**Fig. 8.**
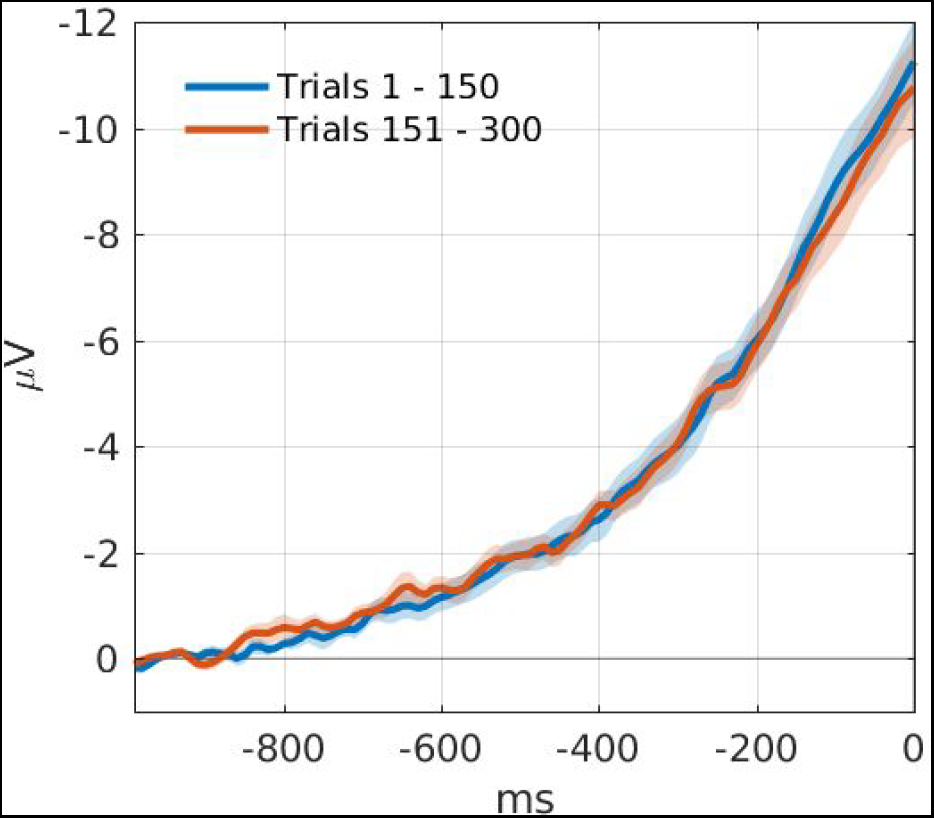
Change of RP waveform in the course of the feedback stage. Grand average RPs in channel Cz, computed for the first (blue) and the second (red) half of trials, respectively. Standard error is shown as a shaded area. Baseline correction was in the interval −1000 to −900).

**Fig. 9.**
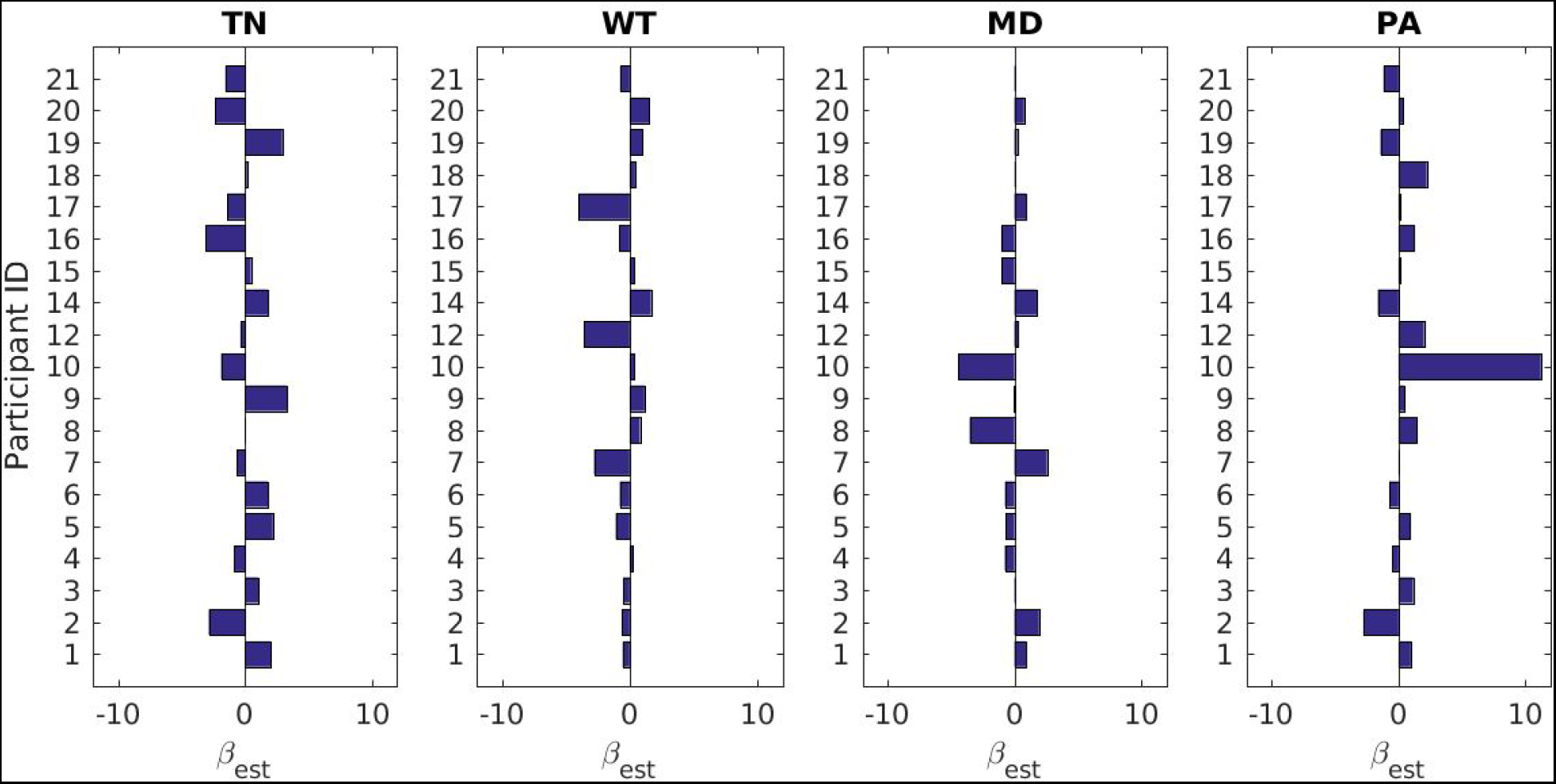
Coefficient estimates from the linear regression. Bar plots show, for each participant, coefficient estimates of the explanatory variables trial number (TN), waiting time (WT), movement duration (MD) and peak acceleration (PA), obtained from a linear regression on the feedback scores.

Next, we examined whether the RP was modulated by either of the three movement characteristics waiting time (WT), movement duration (MD) and peak acceleration (PA). A visual comparison of RPs averaged according to a median split of the three measures of movement characteristics showed only minor differences (Fig. 10), as compared to the inherent variability of RP waveforms (Fig. 6A). There is an apparent small difference in early time periods between short and long waiting times (Fig. 10A) that is not detected as significant in our regression analysis. Our data do not allow us to tell whether this is a spurious effect because any testing of this time period would be post-hoc. Two-sided t-tests on the regression coefficient estimates obtained for each participant (Fig. 9) showed that they were not significantly different from zero (WT: t_(18)_=−1.121, p=0.277; MD: t_(18)_=−0.373, p=0.713; PA: t_(18)_=1.114, p=0.279). Bayes factors for all three variables (WT: BF_01_=2.432; MD: BF_01_=3.955; PA: BF_01_=2.448) furthermore show that the data are more likely under the null hypothesis and indicate a moderate evidence for absence of an effect. Thus, these results suggest the absence of a relationship between RPs and the range of movement parameter variation observed in this study. Please note that the effects of these variables are not of interest for our core research question because they are (i) not used by the participants to improve their scores and (ii) reflect physical changes in the movements.

**Fig. 10.**
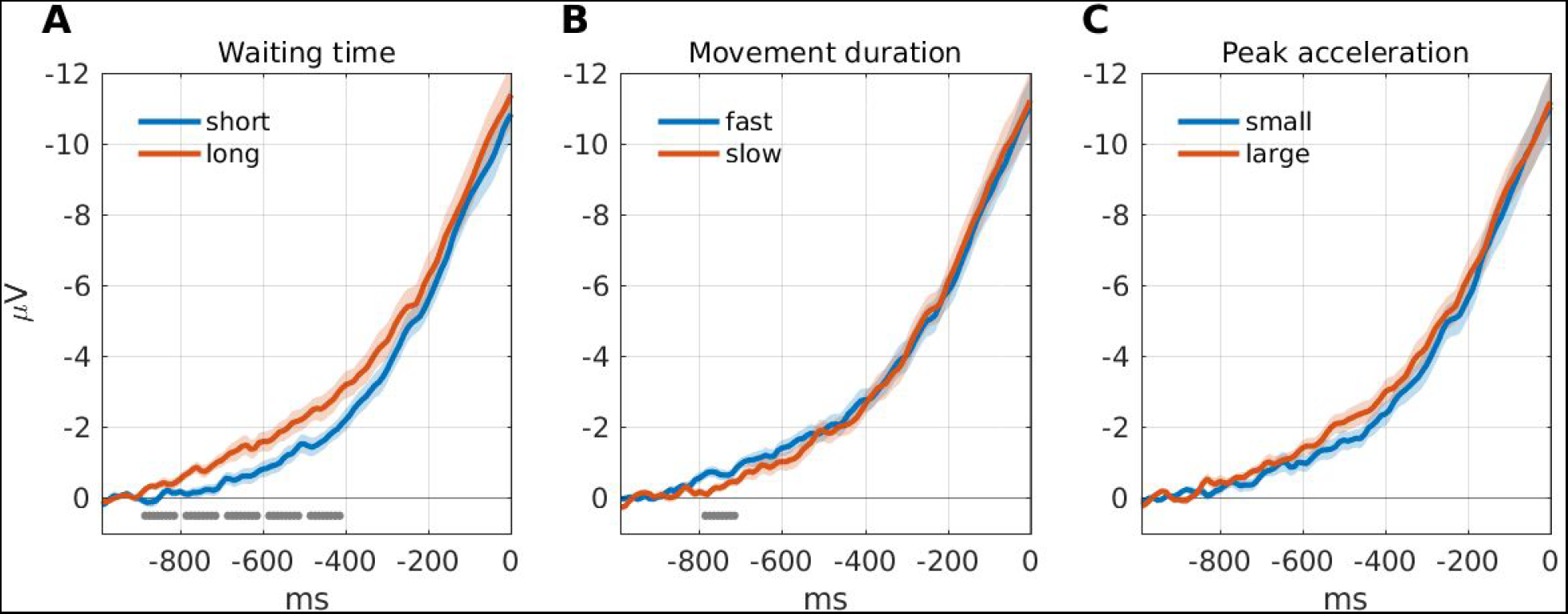
Modulation of RPs by movement characteristics (channel Cz). We additionally checked whether basic spontaneous movement characteristics included in the model as effects of no interest modulated the RP waveforms (as reported in previous literature). For each participant individually, and for the movement characteristic measures (A) waiting time, (B) movement duration, and (C) peak acceleration, two average RPs were generated each using half of the trials (according to a median split of the corresponding measure). The curves show grand averages across participants. Blue and red traces show the average RP for the shorter and longer half of waiting time (A), for the faster and slower half of movements (B), and for the smaller and larger half of peak accelerations (C), respectively. Standard error is shown as a shaded area. Baseline correction was in the interval −1000 to −900. Gray bars indicate in which consecutive, non-overlapping 100 ms windows a paired t-test showed a significant (p<0.05) difference of RP averages. The small differences in the waveforms in A and B did not affect the EEG classifier (and therefore the feedback scores), which takes into account the RP waveform across all selected channels.

### 3.4. Self-assessment of task

We used a questionnaire after the *feedback* stage to assess participants’ experiences and strategies (Fig. 11). When asked to rate how much they felt they could influence scores, the most frequent rating was 3 (“average”). When asked to rate how difficult it was to find a strategy that had an effect on the scores, the most frequent rating1 were 1 and 2 (“very hard” and “rather hard”). Participants were also asked to describe in written form the used strategies and whether they were successful. Among the strategies reported as successful, some participants named a shift of attention towards/away from the movement (2), using relaxation (2), changing the speed (5) or the force (2) of the movement, altering waiting time (2), and involving emotion (2). For detailed reports of participants’ answers, please see Tab. S1 in the Supplementary Material.

**Fig. 11.**
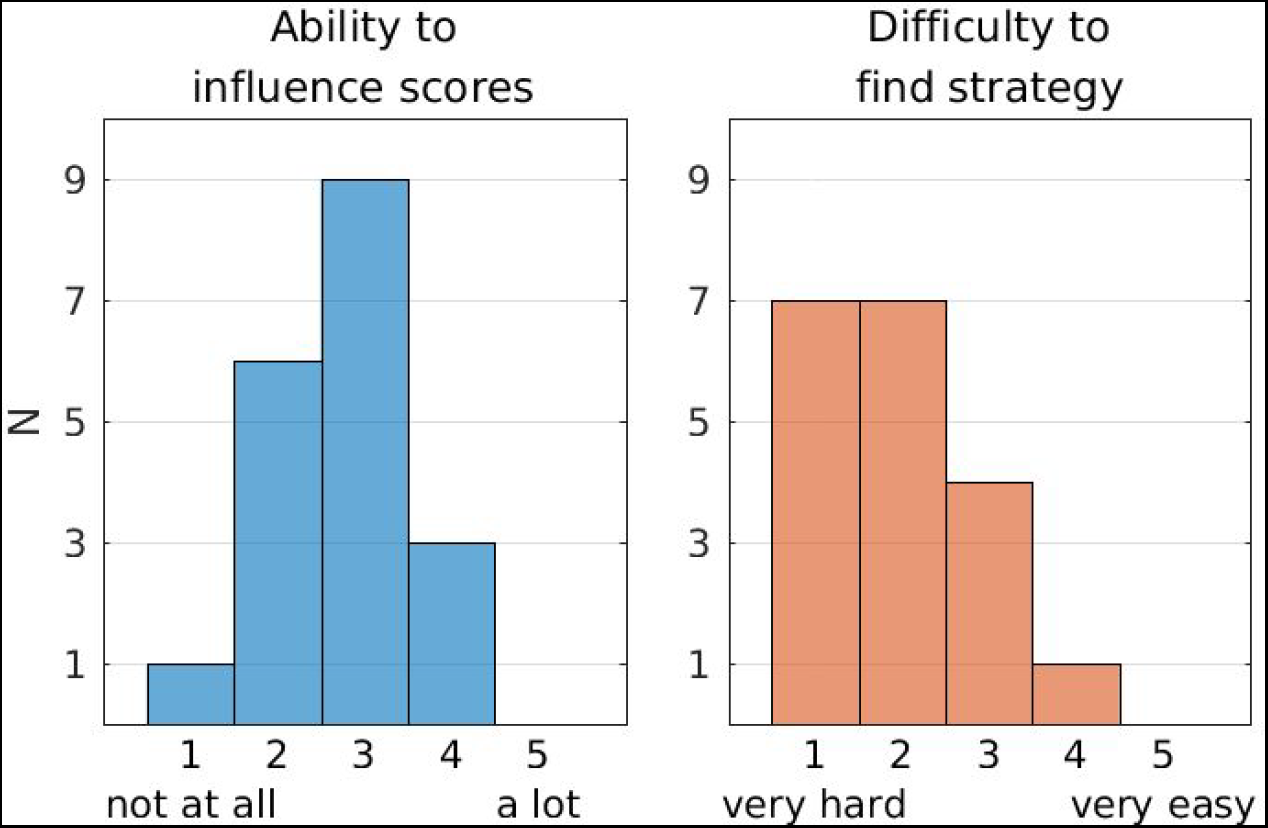
Likert scale rating in questionnaire of subjective experience. Histograms show the number of participants rating question 1 (“Overall, how much did you feel you could influence the scores shown on screen?”) on a Likert scale from 1 (not at all) to 5 (a lot), and question 2 (“How hard/easy was it to find a strategy that had an effect on the scores?”) on a Likert scale from 1 (very hard) to 5 (very easy).

## 4. Discussion

Can people learn to suppress their readiness potential? We tested this possibility in a neurofeedback experiment. Participants performed self-paced pedal presses in single trials and after each pedal press they were provided with a feedback score that reflected the magnitude of the RP preceding that movement.

To extract the scores from RPs, we employed a machine learning approach: We used data acquired during a preparatory stage to train a classifier to distinguish EEG segments preceding movements from the idle period before onset of the trial. By extracting spatio-temporal features from the EEG, the classifier learned both the spatial distribution of the RP across channels, and the characteristics of its waveform. Thus, when the classifier was applied to an EEG segment preceding a movement onset during the feedback stage, the resulting classifier output -- and thereby the feedback score -- was a continuous indicator of the degree to which an RP was present in that segment (Fig. 6A).

Participants were challenged to find a way to perform self-paced movements with small RPs, and were instructed to keep using and extending any potentially effective strategies. If they were successful, this would be reflected in a gradual decrease of feedback scores in the course of the 300 trials of the feedback stage. However, we found no evidence for such a decrease (Fig. 7), suggesting that participants were not able to find or train a successful strategy.

This finding does not rule out the possibility that participants were able to occasionally modulate their RPs. Possibly some weaker potential effects of their strategies went unnoticed and were thus not further explored. One way to test this is to examine the relationship between feedback scores and the movement characteristics that participants were able to modulate and that we could measure in every trial: How long participants waited from trial start until initiating the movement (waiting time), how fast they executed the movement (movement duration), and how much force they applied to the movement, as reflected by the peak acceleration during movement execution. Although waiting time and movement duration slightly modulated the shape of the RP (Fig. 10), we found no evidence for an effect of either of the three movement parameters on the feedback score.

The failure to deliberately suppress the RP does not reflect a fundamental impossibility that small RPs occur. Our data clearly show that many RPs recorded during the feedback stage were remarkably small in size: One out of five movements were preceded by RPs with very late onsets of only a few 100 ms, had amplitudes more than 50% smaller as compared to the average, and a substantially more confined spatial distribution (Fig. 6A, light color code). These small RPs occurred with a fairly constant rate throughout the feedback stage (Fig. 7, light color code). We cannot fully exclude the possibility that the small amplitude of these RPs were somewhat influenced by mental strategies, however if so then participants failed to notice or systematically exploit the effects of these strategies.

When asked how difficult it was to find a successful strategy, most participants (14 out of 19) reported it was hard or very hard (Fig. 11). This self-assessment is in agreement with our finding that scores did not decrease over the course of the feedback stage. Interestingly, there appears to be some kind of “illusion of control”: When asked to rate their general ability to influence scores, participants’ ratings peaked at the midpoint of the scale, reflecting a “moderate” perceived ability to influence scores. In the absence of true control over RP scores this could suggest alternative interpretations. Possibly participants were biased to remember more of those trials in which an intended strategy happened to coincide with a purely random low score, and less those in which the effect was contrary (Thomson et al., 1998). Alternatively, people often tend to choose scores in the middle of Likert scales (which is known as the *central tendency bias*), particularly when they are unsure about their answer (Nadler et al., 2015). Thus, the predominance of central ratings in this question could be interpreted as participants meaning “I don’t know”, rather than as reporting a perceived ability to influence scores. Finally, it is worth noting that we cannot exclude the possibility that participants were effectively able to slightly modulate their RPs, for instance by employing strategies based on attention (Keller & Heckhausen, 1990) which we did not measure and therefore could not test. However, even if that were the case, this modulation effect was too small for participants to train it or to sustain it over longer periods.

At first sight, our data seem to differ from previous findings that paralyzed patients can learn to self-regulate their slow cortical potentials by means of real-time visual feedback (Kübler et al., 1999; Kübler et al., 2001; Neumann et al., 2004; Birbaumer 1999). This raises the question why participants in our study weren’t able to use comparable mechanisms to suppress their RPs. We consider two potential reasons: First, SCPs investigated in those studies reflect changes in cortical polarization that occur *spontaneously* in the ongoing EEG. In contrast, RPs are defined as time-locked to the onset of a voluntary movement, and it has been recently debated whether they occur in the absence of voluntary action (Travers et al., 2020). Thus, it is possible that the event-related nature of RPs impedes their conscious self-regulation by the mechanisms through which other slow cortical potentials are influenced. Second, paralyzed patients learn and train the task of SCP self-regulation in multiple sessions over the course of several weeks or months (Kübler et al., 1999; Kübler et al., 2001; Neumann et al., 2004). While such an approach would be prohibitively expensive for our purposes, we cannot exclude that learning our task might be possible if participants were to be provided with much more time. Finally, it is worth noting that we deliberately did not provide participants with any specific instructions as to how they could achieve the task of suppressing their RPs. We abstained from doing so because we did not make specific assumptions about whether or how this task was possible. Thus, we aimed to test whether a trial-and-error approach was sufficient for participants to find a successful strategy, without introducing a bias on potential strategies. It is however possible that providing specific instructions for mental strategies might have facilitated participants to identify and train a successful strategy.

Our data confirm and expand findings from a recent study, where stop signals were elicited in real-time upon detection of RPs while participants were performing self-initiated movements (Schultze-Kraft et al., 2016). In one condition, participants were instructed to “move unpredictably” so as to not cause stop signals. However, the shape of the RP remained unchanged and stop signals thus continued to be elicited, suggesting that participants were unsuccessful in reducing or suppressing their RPs. In that study, in order to avoid stop signals being triggered by noise in the EEG, they were only elicited if the magnitude of an RP was above a certain threshold. Thus, those stop signals can be considered a *binary* feedback of the RP, since they were triggered by large but not by small RPs. In contrast, in the current study the feedback of the RP was *continuous*: in every trial, a feedback score was shown that directly reflected RP magnitude on a continuous scale. The trial-by-trial feedback in our study thus provided considerably more information about the RP to the participant, compared to the binary stop signals used by Schultze-Kraft and colleagues (2016). However, the data of both studies suggest that the inability of participants to exert control over their RPs does not depend on the type or scale of the provided feedback.

Our main finding that participants were not able to consciously suppress or modulate their RPs suggests that the RP is a signal over which people cannot exert conscious control, and thus that it is an “*involuntary* precursor signal of voluntary action”. Please note, however, that our data remain silent as to whether the RP is a *causal* precursor signal of voluntary action, as has been the traditional account of the RP (Libet et al., 1983; Libet 1985). Alternative accounts suggest that the RP reflects the leaky integration of spontaneous fluctuations in a drift-diffusion process (Schurger et al., 2012; Schurger, 2018), and that spontaneous movements occur when the accumulation of autocorrelated noise reaches a threshold, with either the output (Schurger et al., 2012) or the input (Schurger, 2018) of this accumulation giving rise to the shape of the RP.

The accumulation-to-bound model makes several predictions relevant for the interpretation of our data. First, the RP-as-input model (Schurger, 2018) predicts that the shape of the RP is influenced by the delay between trial start and movement onset, i.e. the waiting time. Indeed, visual inspection of our data show a slight modulation of RP waveform in channel Cz by waiting time (Fig. 10A), compatible with the report by Schurger (2018, Fig. 6). Note, however, that this modulation is not detected as significant in our regression analysis on the feedback scores, where the Bayes Factor supports the absence of an effect. This is possibly because our EEG classifier was trained in a more robust fashion on changes in RP across all channels selected in the training data. Thus, it is unclear if this effect is spurious. Second, in the accumulator model framework people could potentially exert influence over their RPs, e.g. by modulating parameters such as drift rate or threshold, as long as these were in turn to change the shape of the RP. Nonetheless, if this accumulation is necessary for voluntary movements then such movements would necessarily be preceded by an RP. Our data show that participants seem unable to affect the amplitude of the RP, even when explicitly trying to do so.

In sum, we performed a neurofeedback experiment to test whether people are able to suppress their readiness potential. We found no evidence for the ability of participants to consciously suppress their RPs. Our findings thus suggest that the readiness potential is an involuntary precursor signal of voluntary action over which people cannot exert conscious control.

## Acknowledgements

This work was supported by a joint grant from the John Templeton Foundation and the Fetzer Institute (to MSK, JDH). The opinions expressed in this publication are those of the author(s) and do not necessarily reflect the views of the John Templeton Foundation or the Fetzer Institute. Furthermore, this work was supported by the Excellence Cluster “Science of Intelligence” (DFG SFB 904/1), the Collaborative Research Center “Volition and Cognitive Control” (DFG EXC 2002) and the Max Planck School of Cognition (to JDH, JS). Finally, this work was supported by Erasmus+ traineeship program (to TB).

## Supplementary Material

### Participants’ self-reports on questionnaire about strategies

**Table S1.** Participants’ self-reports on questionnaire about strategies. Participants were asked
to write down (i) the different strategies that they used during the feedback stage, (ii) whether they worked, and if so (iii) whether they were trainable. The table shows all reported strategies of single participants in condensed form. Participants’ written accounts of how well each strategy worked were classified into three rating categories (right column): accounts such as “unsuccessful” or “did not work” are denoted with a 0, accounts such as “seemed to work” or “worked sometimes” are denoted with a +, accounts such as “successful” or “worked well”, or those reported as trainable, are denoted with ++.

| Participant ID | Strategy | Rating |
| --- | --- | --- |
| 1 | Adjust breathing | 0 |
|  | Focus on the impulse to press | + |
|  | Shift attention to something else | + |
|  | Relax | 0 |
| 2 | Focus on the visual stimulus | 0 |
| 3 | Plan the next movement | 0 |
|  | Ignore the goal | 0 |
|  | Reward myself for good scores | 0 |
| 4 | Relax and clear the mind | + |
|  | Evoke positive emotions | ++ |
|  | Ignore scores | 0 |
| 5 | Relax, don't think about pressing | ++ |
|  | Move faster | 0 |
|  | Think about low numbers | 0 |
| 6 | Clear the mind | + |
|  | Play song in my head and hit on specific beats | 0 |
|  | Delay the first urge to press and press later | 0 |
| 7 | Don't think about pressing | + |
|  | Focus on pressing | ++ |
|  | Delay first urge to press and press later | + |
|  | Feel surprised about pressing the pedal | ++ |
| 8 | Slow down movement and decrease force | + |
| 9 | Delay the urge to move and move shortly after | ++ |
|  | Don't think about pressing | + |
|  | Longer waiting time | 0 |
| 10 | Focus on stimulus or task | 0 |
|  | Focus on breathing, clearing the mind | 0 |
|  | Slower and less forceful movements | ++ |
|  | Longer waiting time | 0 |
| 11 | Longer/shorter waiting time | 0 |
|  | Increase/decrease force of movement | 0 |
|  | Interrupting movement | ++ |
| 12 | Relax | 0 |
|  | Slower movement | ++ |
| 13 | Slower movement | ++ |
|  | Decrease force of movement | 0 |
|  | Decrease waiting time | 0 |
| 14 | None specified | 0 |
| 15 | Mind wandering (use spontaneous train of thoughts) | ++ |
|  | Relax | + |
|  | Shifting attention away from task | + |
|  | Using emotions | + |
| 16 | Increase force of movement | ++ |
|  | Shift attention away from task | + |
|  | Increase waiting time | 0 |
|  | Relax | + |
| 17 | Clear mind, relax | + |
|  | Move spontaneously | + |
| 18 | Think about something else | 0 |
|  | Focus on breathing | + |
|  | Focus on replicating the same movement every trial | ++ |
|  | Shift attention away from task | 0 |
|  | Increase movement speed | 0 |
| 19 | Shift attention away from task | + |
| 20 | Waiting time | ++ |
|  | Force of movement | 0 |
|  | Movement speed | ++ |
| 21 | Moaning | + |
|  | Relax | 0 |
|  | Speed of movement | 0 |
| 22 | Delay the first urge to move, move later | + |

